# A comprehensive survey of coronaviral main protease active site diversity in 3D: Identifying and analyzing drug discovery targets in search of broad specificity inhibitors for the next coronavirus pandemic

**DOI:** 10.1101/2023.01.30.526101

**Authors:** Joseph H. Lubin, Samantha G. Martinusen, Christine Zardecki, Cassandra Olivas, Mickayla Bacorn, MaryAgnes Balogun, Ethan W. Slaton, Amy Wu Wu, Sarah Sakeer, Brian P. Hudson, Carl A. Denard, Stephen K. Burley, Sagar D. Khare

## Abstract

Although the rapid development of therapeutic responses to combat SARS-CoV-2 represents a great human achievement, it also demonstrates untapped potential for advanced pandemic preparedness. Cross-species efficacy against multiple human coronaviruses by the main protease (MPro) inhibitor nirmatrelvir raises the question of its breadth of inhibition and our preparedness against future coronaviral threats. Herein, we describe sequence and structural analyses of 346 unique MPro enzymes from all coronaviruses represented in the NCBI Virus database. Cognate substrates of these representative proteases were inferred from their polyprotein sequences. We clustered MPro sequences based on sequence identity and AlphaFold2-predicted structures, showing approximate correspondence with known viral subspecies. Predicted structures of five representative MPros bound to their inferred cognate substrates showed high conservation in protease:substrate interaction modes, with some notable differences. Yeast-based proteolysis assays of the five representatives were able to confirm activity of three on inferred cognate substrates, and demonstrated that of the three, only one was effectively inhibited by nirmatrelvir. Our findings suggest that comprehensive preparedness against future potential coronaviral threats will require continued inhibitor development. Our methods may be applied to candidate coronaviral MPro inhibitors to evaluate in advance the breadth of their inhibition and identify target coronaviruses potentially meriting advanced development of alternative countermeasures.

## Introduction

The COVID-19 pandemic, caused by the betacoronavirus SARS-CoV-2^1^, has for the third time demonstrated the danger posed by animal coronaviruses jumping the species barrier to humans. The current pandemic was preceded by similar betacoronavirus jumps giving rise to SARS and MERS epidemics in 2002 and 2012, respectively. Other zoonotic transmission events that have not spread between humans to produce epidemics are thought to be common^2^, and are not limited to alpha- and betacoronaviruses^3^. Moreover, it appears that many SARS-related coronaviruses are sufficiently similar to human epidemic viruses that small genomic/proteomic sequence changes may turn them into effective pathogens^4^. As coronaviruses are highly recombinogenic, it is conceivable that diverse, currently non-pathogenic viruses may acquire pandemic/epidemic potential. Consequently, it appears highly likely that humanity will again need to address the threat of novel infectious coronaviruses spilling over and causing regional epidemics or, if not halted promptly, pandemics.

Since we do not know which coronavirus might be next to make the jump to humans, it behooves us to evaluate the full diversity of known *coronaviridae* to be more aware of potential threats. We and others have observed that some coronaviral proteins are less variable than others; the spike protein, which is responsible for virion binding and entry to host cells and is the target for the SARS-CoV-2 vaccines, undergoes frequent alterations, including those occurring within the receptor binding domain^5^. The proteases that liberate individual proteins from viral polyproteins after translation are significantly more conserved than the spike protein, particularly in their active sites. The main protease (MPro, nsp5) is responsible for peptide bond hydrolysis at 11 cleavage sites within the polyprotein^6^. Due to the relative stability of its active site and functional importance, SARS-CoV-2 MPro represents an attractive target for inhibitor discovery^7,8^. The degree of conservation, even among different coronaviral species, is such that an inhibitor for the SARS MPro likely would have been effective advance countermeasure against SARS-CoV-2 MPro^9^.

In this work, we develop a pandemic preparedness pipeline by assessing the diversity of MPro active site 3D structures (experimentally-determined and computationally-modeled) across all known coronaviruses. We evaluate the diversity of coronaviral MPro enzymes, including all known genera (*alpha*-, *beta*-, *gamma*-, and *deltacoronaviridae*). We developed a nonredundant set of MPro sequences extracted from the NCBI Virus database^10^ and generated computed structure models (CSMs) of each. We examined their active sites, and selected a set of five representative models based on active site sequences. We generated CSMs of these representatives in complex with their native substrates, inferred from their polyprotein sequences, and compared them to those of experimentally-determined SARS-CoV-2 MPro-substrate complexes. We also performed conservation analyses of active sites and substrates across all known species. Finally, we employed a yeast surface display-based activity assay with our chosen representative MPros to ascertain substrate cleavage and inhibition by nirmatrelvir^8,11^.

## Results

### Assessing MPro sequence diversity

We assessed sequence diversity of the MPro enzymes across all coronaviruses whose genome sequences are available. We assembled a dataset of 346 MPro sequences from the NCBI Virus genome sequence database^10^ (< 99% protein sequence identity) for comparison. This dataset represents the entire breadth of documented coronaviral species. The pairwise sequence comparisons range from <35% sequence identity to 99% (mean: 52%, SD: 20%) (Table S1). As shown in the hierarchically clustered cladogram (Figure 1A), the MPros tend to cluster by *genera* (alpha, beta, gamma, and delta coronaviruses), although there are exceptions. SARS and SARS-CoV-2, both betacoronaviruses, are closely related, but are not the closest known neighbors in the phylogenetic tree constructed from MPro sequences alone. MERS, also a betacoronavirus, is slightly more distant from SARS and SARS-CoV-2 than they are from each other. There is relative proximitybetween these species among the full family of *coronaviridae*.

**Figure 1.**
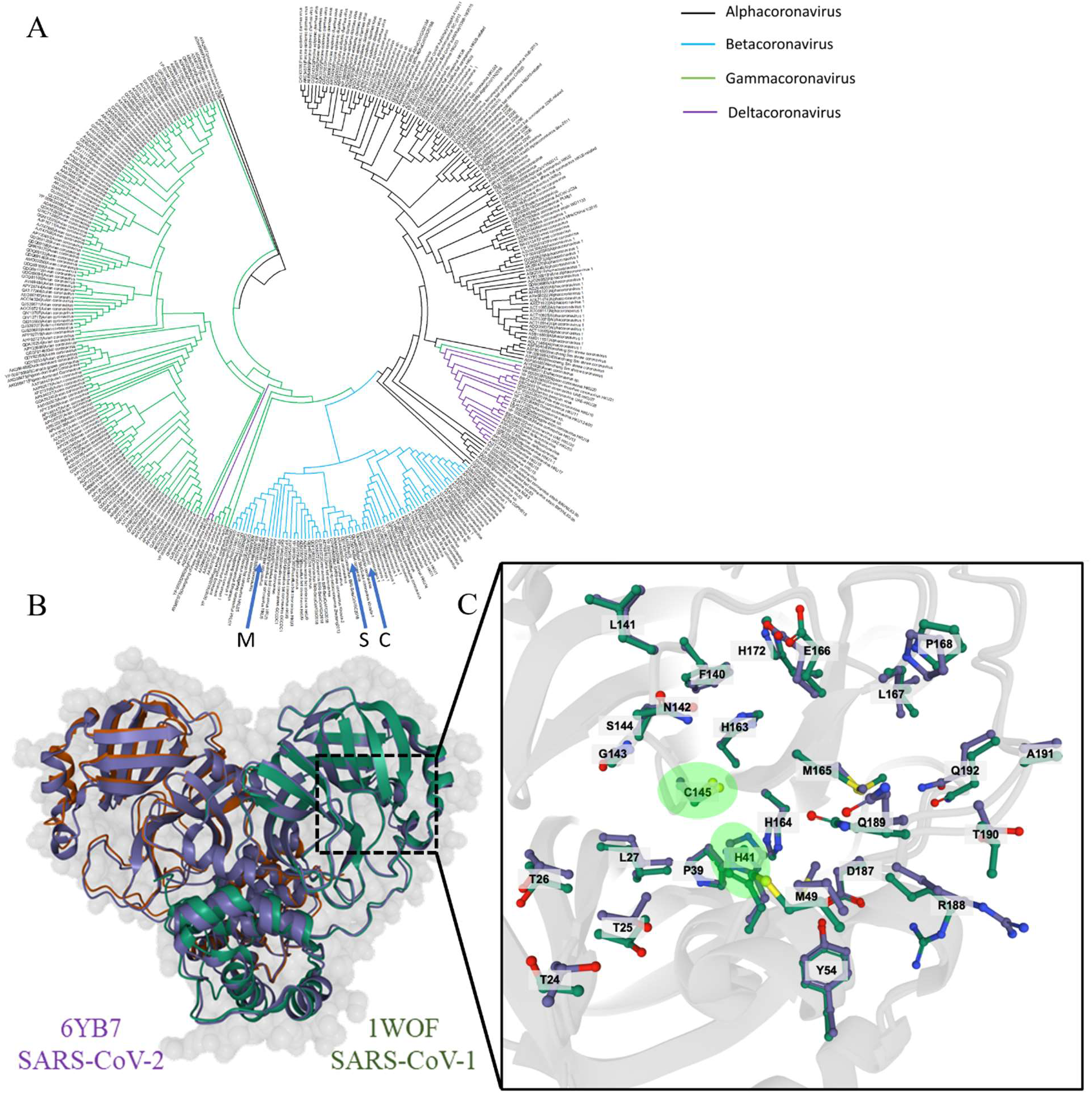
A: Full protease cladogram. Phylogenetic tree organized hierarchically based on full amino acid sequences for known MPro enzymes. Colors reflect coronaviral genera. Blue arrows indicate MERS (M), SARS (S), and SARS-CoV-2 (C). B and C: Structural superposition of SARS (PDB ID 1WOF) and SARS-CoV-2 (PDB ID 6YB7) MPro structures. B: Full MPro homodimer. C: Active site residues. Catalytic residues are highlighted green.

Elucidating the relationship between active site sequence and substrate recognition in the identified representative MPros requires examining molecular structures in 3D, particularly at the active site (Figure 1B and C). However, experimental 3D structures were not available for the vast majority of MPro enzymes represented in the NCBI Virus database. We generated Computed Structural Models (CSMs) of all identified MPros and selected five representatives based on clustering as described in Methods. To provide the highest quality CSMs, we generated both monomeric and dimeric structures of each representative MPro using AlphaFold 2 (AF2). MPro is a symmetric homodimer in SARS-CoV-1, MERS, and SARS-CoV-2, and we reasonably assumed that this is the case for all coronaviral MPros.

### Active site clustering and representative selection

All coronaviral MPro inhibitors target the active sites of these enzymes. For this reason, we focused our analyses on the enzyme active sites. First, we identified residues lining the active sites in known macromolecular crystallography-determined (MX) structures of SARS-CoV-2 MPro, Figure 2A). We then identified homologous residues in the MPro CSMs using Dali^12^. Within the active site-only sequences, diversity is reduced compared to the full MPro sequence (mean sequence identity: 66%, SD: 17%) (Table S2). We expect that this is the result of functional conservation for substrate binding. A second hierarchical clustering, based exclusively on the protease active site sequences (Figure 2B), also results in clusters that correspond approximately to known coronaviral *genera*. The sequences are grouped into five clusters (whereas there are four *genera*), with lineage mixing most notable in Cluster 3, which is why it was considered distinct from the adjacent clusters. The subsequent analyses use these five identified clusters.

**Figure 2.**
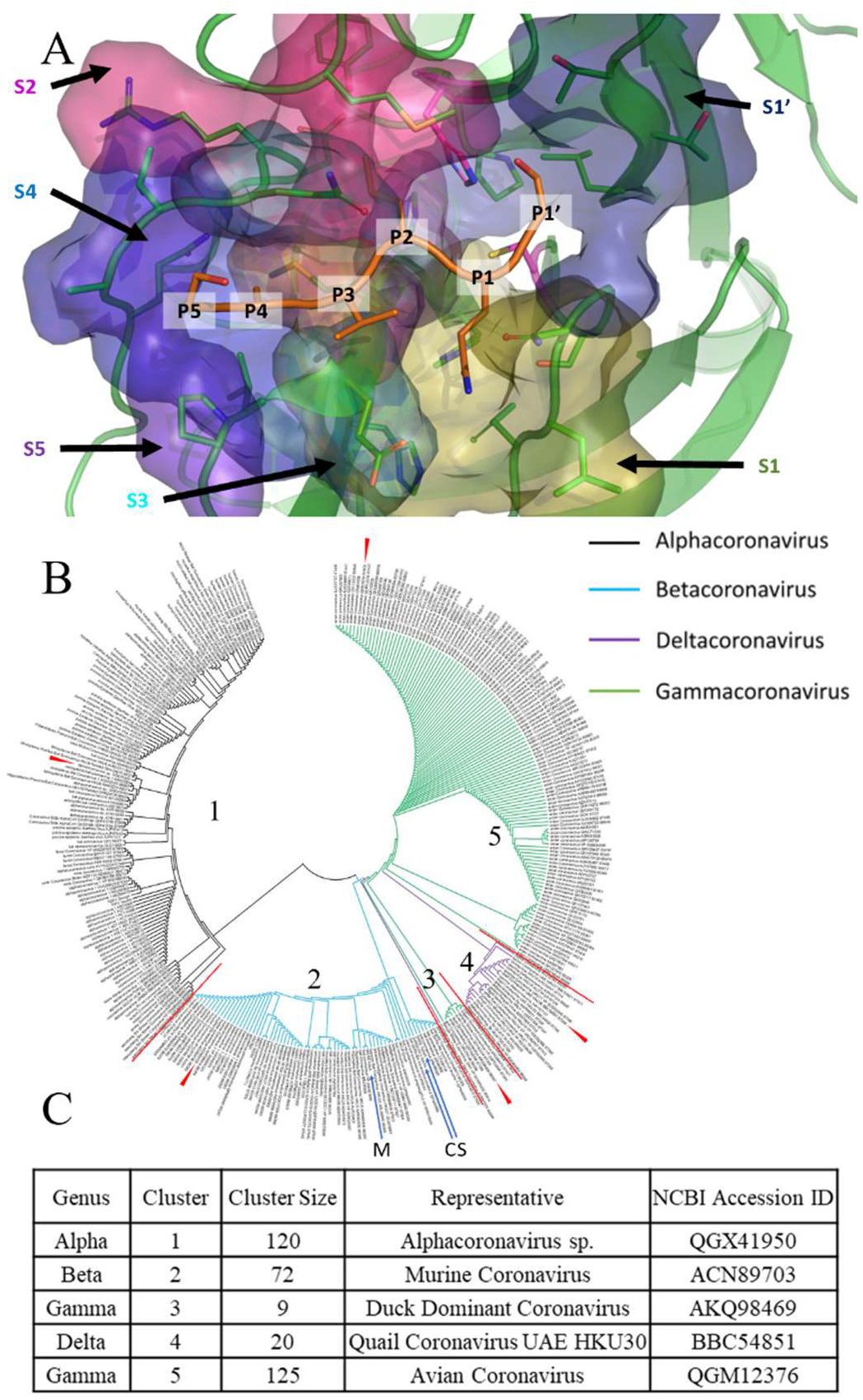
Active site sequence diversity for all coronaviral MPros. A: Sub-pockets of the active site shown on a computed model of SARS-CoV-2 MPro binding a native substrate, ITSAVLQ/SGF. Sub-pockets are numbered according to contacted substrate residue, i.e. sub-pocket S5 contacts substrate residue P5. B: Active site phylogenetic tree organized hierarchically based on MPro active site sequences. Colors reflect coronaviral genera. Blue arrows indicate MERS (M), SARS-CoV-2 (C), and SARS (S). Numbers indicate clusters used for identifying representatives, which are separated by red lines. Red wedges indicate selected representative structures. C: Selected representative models based on protease active sites.

We sought to identify representative CSMs that maximally encapsulate the diversity of known coronavirus MPros against which existing inhibitors might be screened to determine potential breadth of their efficacy. For this analysis, we selected only one CSM from each cluster, shown in Figure 2B and C. However, the hierarchical nature of the clustering allows for rapid distinction of smaller active site sequence clusters for selection of additional representative CSMs, should greater granularity be desired. We selected a single representative of each cluster based on maximum structural similarity of the CSM to all other cluster members, based on an all-vs-all Z-score matrix obtained from Dali. We then compared our representative CSMs with experimentally-determined SARS-CoV-2 MPro active site structure in complex with substrates.

### Modeling Enzyme-Substrate Complexes

The activity of an effective inhibitor often depends on mimicking the substrate of its target enzyme, thus we sought to understand the diversity of protease-substrate interfaces among *coronaviridae* at the molecular level. To benchmark our approach for MPro-substrate complex modeling in homologous MPro enzymes, we sought to recapitulate the MX structures of SARS-CoV-2 MPro-substrate complexes, using the AF2-generated apo structure and a Rosetta-based minimization protocol to generate CSMs for SARS-CoV-2 and its native substrates. Zhao *et al.*^13^ used a SARS-CoV-2 MPro H41A catalytic knockout variant to determine the crystallographic structures of SARS-CoV-2 MPro-substrate complexes. They observed consistent placement of substrate residues P5 (five residues towards the N-terminus from the cleavage site) through P1’ (the residue at the C-terminal side of the cleavage site), with greater positional diversity at P2’ and beyond, and were able to describe a number of the interactions underlying the specificity contributions of sites S4 (the subpocket of the protease interacting with substrate residue P4), S2, S1, and S1’. Residues P5 (A, E, F, G, H, R, S, T, V, Y in SARS-CoV-2 substrates) and P3 (K, M, R, T, V in SARS-CoV-2 substrates) are more solvent-exposed, and thus have fewer constraints on accepted amino acids, though favorable sidechain interactions are possible, particularly for P3. Zhao *et al*. noted H-bonds with Q189 and N142. S4 is a spatially constrained pocket, favoring smaller P4 residues (A, P, T, V). S2 is highly hydrophobic, producing a similar pattern for P2 (i.e., preference for F, L, V). P1 is exclusively Q, because it is able to form three H-bonds, involving F140, N142 (with a bridging water molecule), and H163. S1’ is also a shallower pocket, favoring smaller P1’ amino acid residues (A, N, S).

Our CSMs (Figure 3 and Figure 4A, structures provided in SI) were a close match to those of Zhao *et al.,* including similar backbone (Figure 3A), the consistency of P5-P1’ placement (Figure 3B), and many positions of active site sidechains (Figure 3C), including the catalytic dyad. We also observed some differences in H-bonding patterns (Figure 3D-E). In all CSMs, our placement of E166 is rotated compared to the experimental models, precluding a bond with H172, but still forming bonds with P1, and when applicable, P3. In all cases, however, we recapitulated the network connecting P1 to N142 and H163, and the backbone H-bond with Q189 was recapitulated. In substrates with R at P3 (nsp14 and nsp15), our CSMs placed the sidechain towards E166, whereas the experimental structures indicate bonds with N142 and Q189. The high agreement of our CSMs of SARS-CoV-2 MPro-substrate complexes with corresponding experimentally-determined structures allows us to proceed with modeling holoenzyme complexes of the MPro representatives to examine their active site interactions. Together with the collected assessment of active site and substrate diversity, these understandings of the mechanisms underlying specificity can be used to rationally guide searches for broad inhibitors.

**Figure 3.**
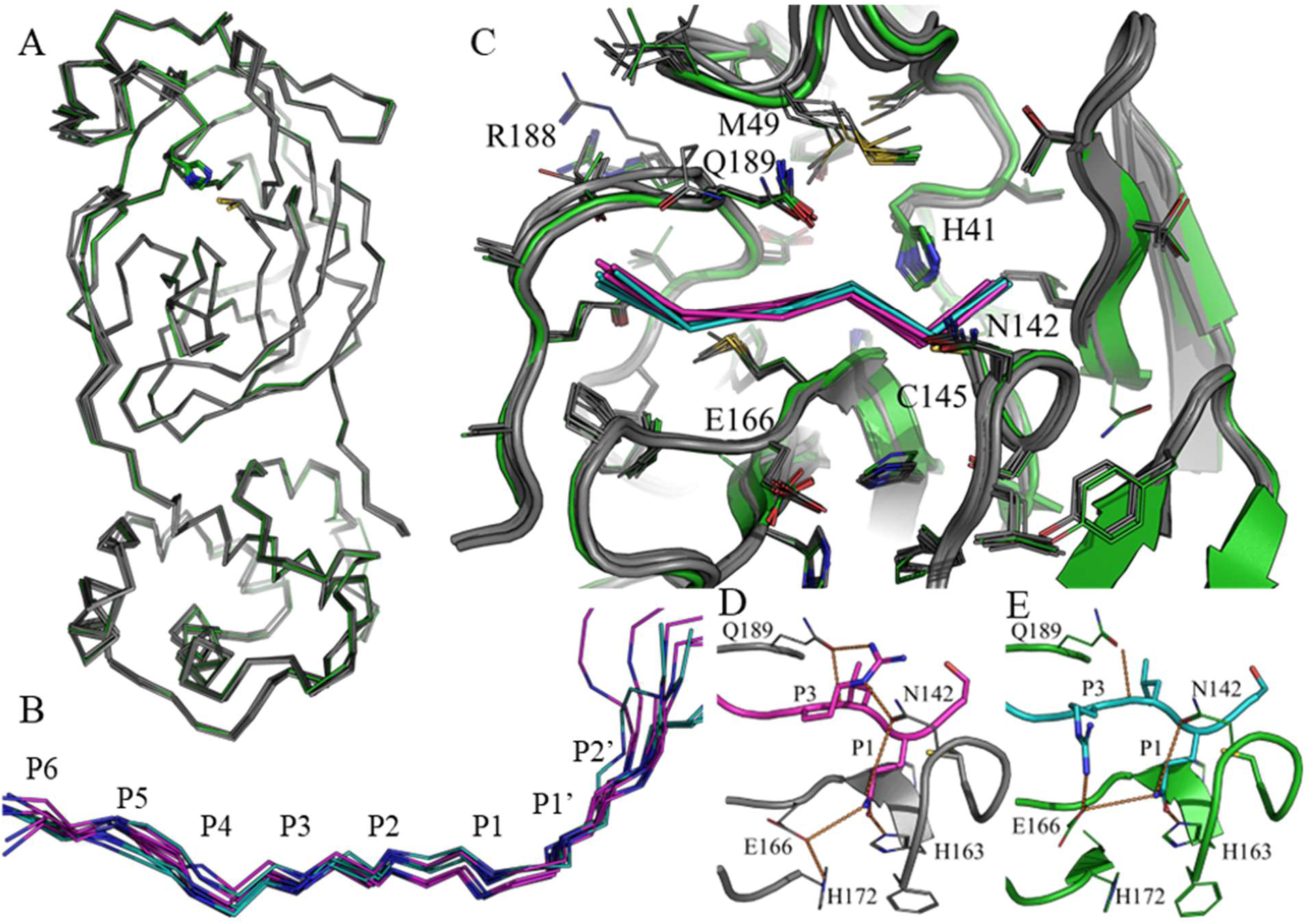
AF2+Rosetta modeling of SARS-CoV-2 holoenzymes shows high agreement with experimentally-determined structures. Experimental structures (7DVP, 7DVW, 7DVX, 7DVY, 7DW6, and 7DW0) are shown in gray (protease) and magenta (peptide); CSMs are shown in green (protease) and cyan (peptide). H-bonds common to both models are shown in orange, for experimental-only in yellow, and for CSM-only in brown. A: Protease backbone alignment. B: Peptide backbone placement when proteases are aligned. C: Active site focus, showing protease side chains. D-E: nsp14 substrate with sidechains and H-bonds shown. D: experimentally-determined structure. E: Computed structure model.

**Figure 4.**
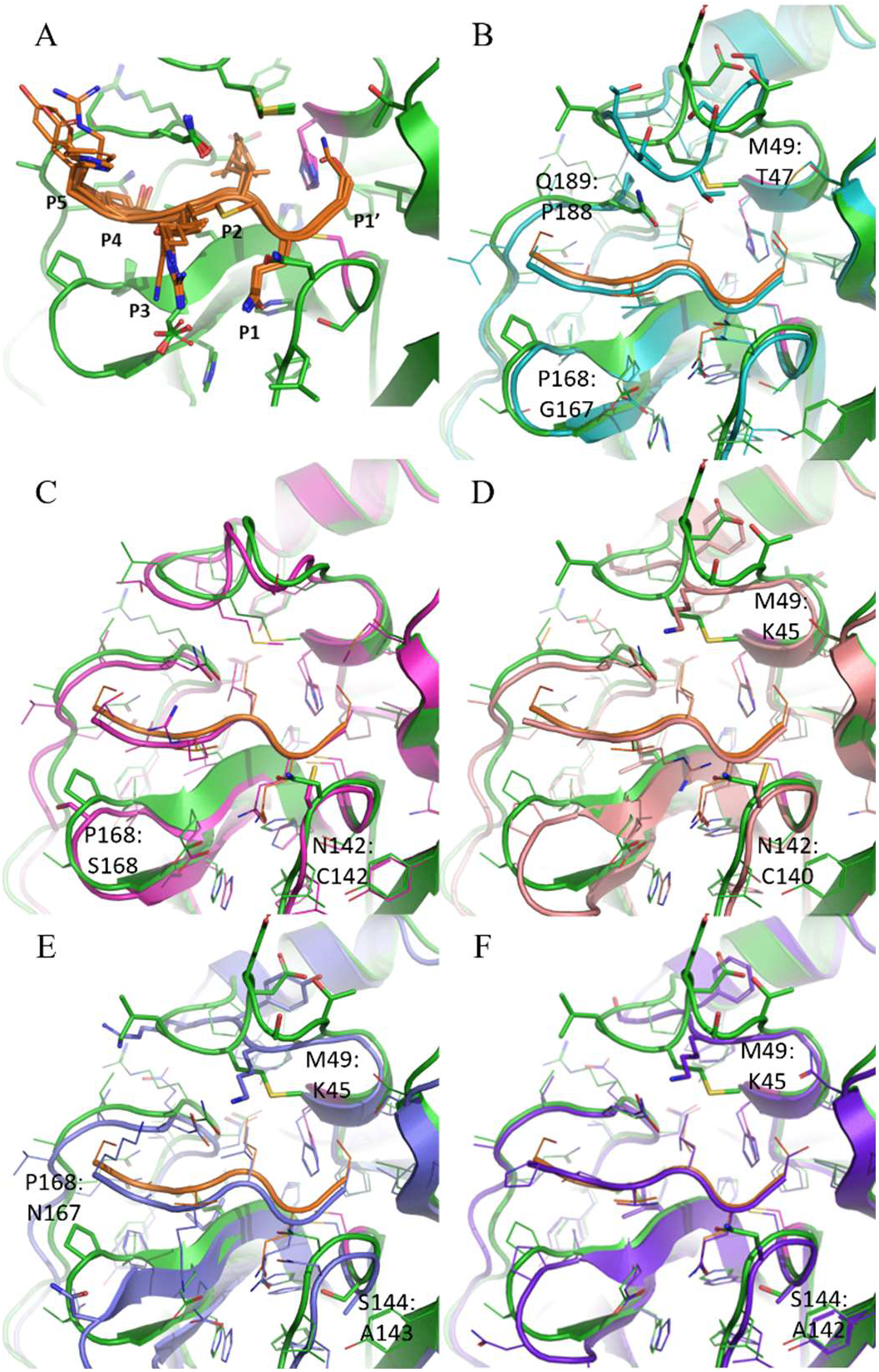
MPro holoenzyme active site structures. A: SARS-CoV-2 MPro with all 11 native substrates. SARS-CoV-2 MPro is in green, except for catalytic H41 and C141, which are in magenta, and substrates are in orange. B-F: Comparison of representative structures to SARS-CoV-2 MPro with thicker sticks showing active site residue changes and side chains on alternately structured loops. SARS-CoV-2 MPro is shown in green. B: MPro 1; C: MPro 2; D: MPro 3; E: MPro 4; F: MPro 5. Residue differences are noted as [SARS-CoV-2 MPro]:[Representative MPro].

We inferred the native substrates of our representatives from their polyprotein sequences (see Methods). Applying the Rosetta-based protocol benchmarked for SARS-CoV-2 MPro-substrate complexes, we generated similar holoenzyme CSMs for each cluster representative, using available substrate sequences identified from polyprotein sequence alignment (Figure 4B-F, Table S3, structures in SI). As our sequence analyses suggested, active sites are highly similar between the clusters. Compared to SARS-CoV-2, the representative of Cluster 1 (Figure 4B) has a different loop structure around residue 46, which appears to be the most variable region near the active site among selected representatives. This structural difference may impact S2, replacing M49 with threonine. Additionally, the substitution of Q189 for a proline may eliminate hydrogen bonding with P3. The most notable difference with the Cluster 2 representative (Figure 4C), which is otherwise structurally similar to SARS-CoV-2 (which is a member of Cluster 2), is the exchange of N142 for a cysteine, which may reduce the stringency for P1 for Q. The Cluster 3 representative (Figure 4D) has a similar cysteine placement. The loop including residue 46 is altered in this structure as well, placing a lysine roughly in the same position as M49 in the Cluster 1 representative structure. The structure of the loop of the Cluster 4 representative (Figure 4E) is similar to that of the Cluster 3 representative, which also has a lysine residue at position 49. The other notable difference is an asparagine instead of proline at position 168, which may form hydrogen bonds with P3 and/or P5 amino acids. The Cluster 5 representative (Figure 4F) has a loop structure similar to those of Clusters 3 and 4. N142 is replaced by an alanine in this case. Thus, while all examined structures have a high degree of active site similarity, the active site pockets of representative MPros have a number of differences that may impact inhibitor binding.

### Active site and substrate sequence comparisons using structure-based sequence alignment

After comparing the active sites of representative structures in atomic detail, we sought to perform a broader analysis of diversity across all collected species. Based on structure-based sequence alignments of all 346 MPro homologs, we compared the prevalence and conservation of homologous active site residues (Figure 5). Figure 5A includes active site residue frequencies across the full dataset (active site residue frequencies for individual clusters are in Tables S4.1-S4.6). We calculated the sequence entropy at each site and found high conservativity (ranging from 0.00 to 1.89, noting that a fully diverse amino acid distribution has an entropy of 3.00). Indeed, members of the catalytic dyad are by no means the only active site residues that are totally or strongly conserved across all known species; nearly all active site residues exhibit some conservation. Many active site amino acid positions were highly conserved across the 346 MPro sequences in our dataset: 27, 39, 140, 143, 163, 166, 172, 187, 192. These are not the catalytic residues, and yet these sites exhibit near-total conservation across all species. Many more exhibited common trends among variants, such as a consistent tendency for hydrophobic residues at sites 141, 165, 167, and 191, aromatics (W or Y) at 54, a small residue at site 144, and a hydrogen bond donor (H or Q) at 164. The sub-pockets with the highest conservativity (i.e. lowest average sequence entropy) are, in order, S1, S3, S4, S1’, S2, S5. Some residues are shared among multiple sites, so their specificity contributions may involve multi-body interaction networks, as opposed to single-residue interactions.

**Figure 5.**
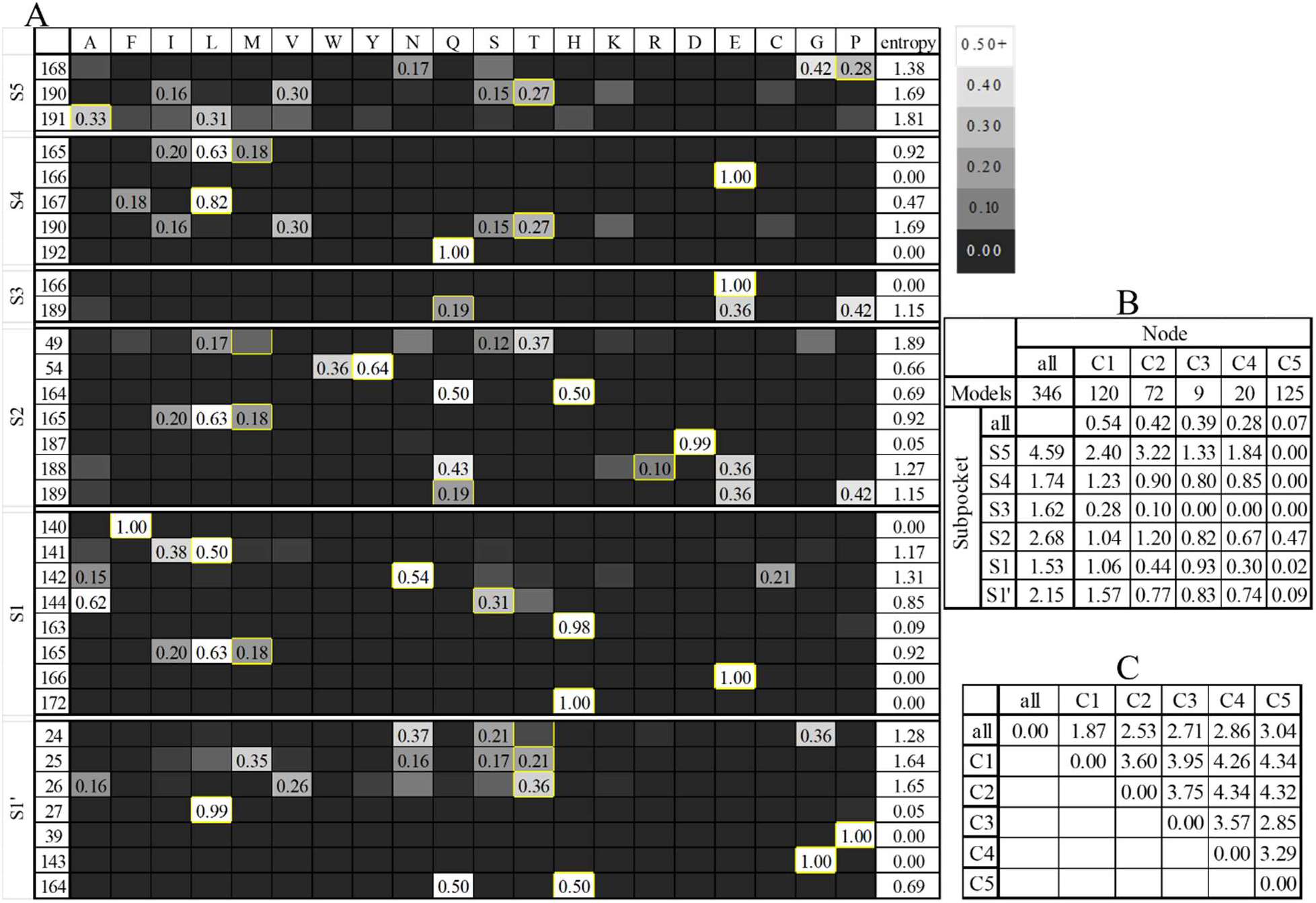
MPro active site sequence diversity. A: Positional residue frequencies in MPro active sites. SARS-CoV-2 residues at each site are denoted with a yellow perimeter. B: Normalized sequence entropy totals for each active site sub-pocket by cluster. (The ‘all’ column is the sums from A.) All values are divided by the number of residues in the subpocket and normalized to the all-clusters all-subpockets entropy, which is defined as 1. C: Frobenius distances between cluster site frequency matrices for all active site residues, not separated by sub-pocket.

We performed a comparison of the total entropies for each sub-pocket, both for the full set of MPros and separately by cluster (Figure 5B, Table S5). Pocket entropy may be used as a metric for overall conservation in each sub-pocket across different clusters. Highly conserved sites may represent potentially more desirable inhibitor binding targets. Despite being the largest group, Cluster 5 has the lowest overall entropy, suggesting that the *gammacoronaviridae* active sites are particularly invariant. In contrast, Cluster 1 has the highest entropy, suggesting a greater challenge for broad inhibition within the *alphacoronaviridae*, which include several human pathogens.

Comparison between the active site residue frequency tables also allows for calculation of Frobenius distance (measure of the distance between two points on the Stiefel manifold)^14^ between different groups (Figure 5C for the full set and by individual sub-pocket in Tables S6.1-S6.6), indicating the degree of dissimilarity between clusters. More proximal clusters may have a higher probability of binding common inhibitors.

MPro active sites coevolved with their cleavage sites within the polyprotein. An amino acid change within a given MPro active site would render the new virus unable to infect its host unless either the mutation does not interdict cleavage, or simultaneous compensatory mutations occur in affected cleavage sites to restore activity. During productive viral infection, MPro must cleave a total of eleven sites within the polyprotein, thus the enzyme and the eleven cleavage sites (listed in Table S7) are collectively subject to evolutionary selection pressure favoring conservation. By aligning the polyprotein sequences from each member of our dataset with the SARS-CoV-2 polyprotein sequence (Table S8), we identified homologous cleavage sites (Table S9). Figure 6 shows the diversity of the identified cleavage sequences. Figure 6A includes all identified cleavage sequences, while residue frequencies are tabulated filtered by cluster in Tables S10.1.1-S10.1-5, by cleavage site in Tables S10.2.1-S7.2.11, and by both in Tables S10.3.1-S10.3.29. Conservation was detected at all cleavage site positions between P4 and P1’, with order of highest conservativity (*i.e.*, lowest sequence average entropy) being P1, P2, P1’, P4, P3, P5. The P1 residue was 99% Q demonstrating a nearly absolute preference for this amino acid. P2 was largely hydrophobic aliphatic sidechains L and V. Notably, the order of conservativity of substrate sites differs from that of active site sub-pockets, most prominently at S3/P3, where the sub-pocket residues are all shared with other sub-pockets, but the substrate sidechain is oriented away from the protease. Mapping these conserved substrate positions onto the MPro active site serves to identify the S1 and S2 subsites as favorable binding regions for a broadly active inhibitor which may maintain potency against diverse MPros.

**Figure 6.**
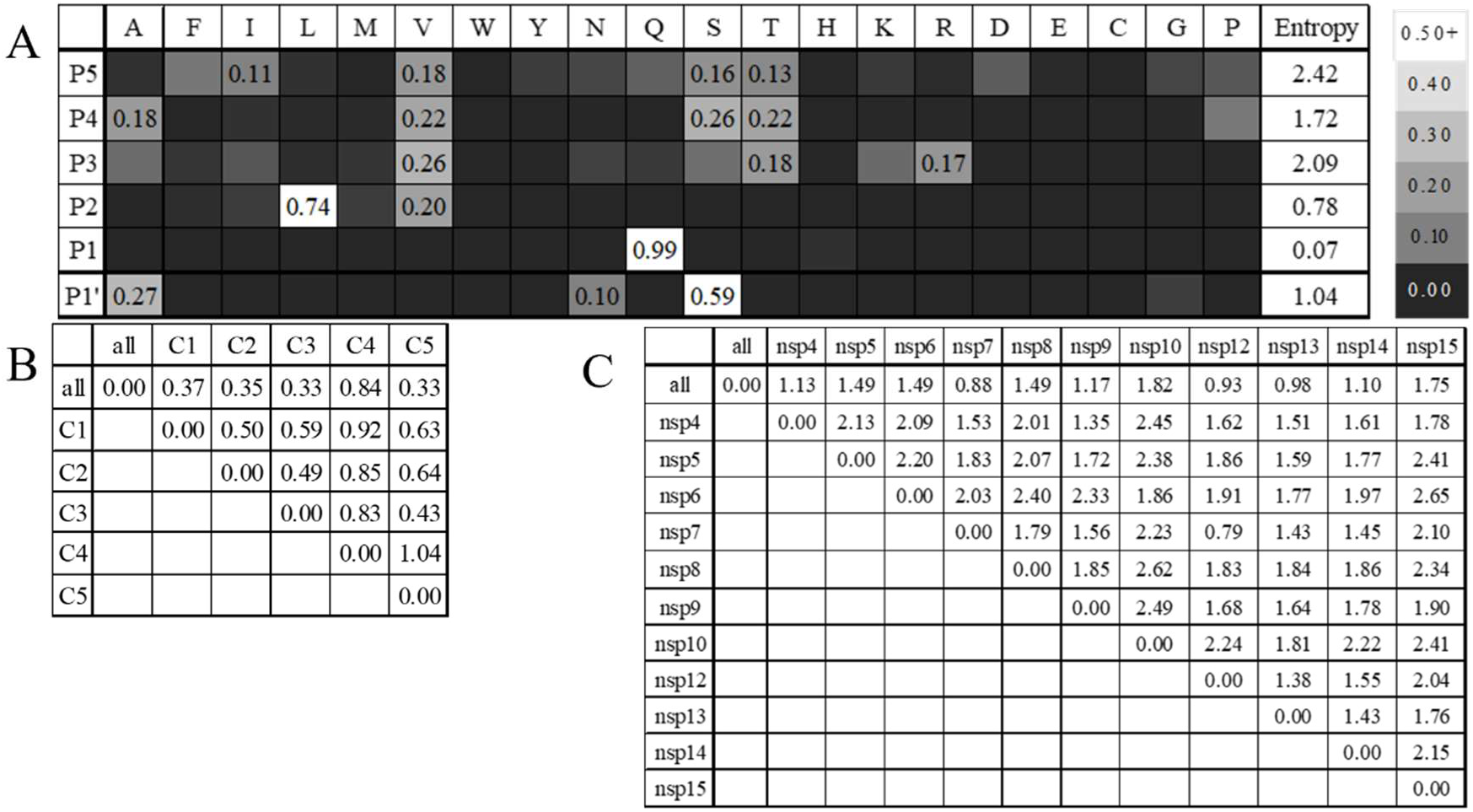
MPro substrate sequence diversity. A: Positional residue frequencies in MPro substrates. Cleavage occurs between P1 and P1 ‘. Increasing numbers indicate distance from the cleavage site in the N- and C- terminal directions, respectively. B-C: Frobenius distances between frequency matrices. Listed cleavage sites indicate the polyprotein member on the N-terminal side of the cleavage site, so nsp4 is the site between nsp4 and nsp5, with sequence SAVLQ/S. B: Substrate sequences by cluster. C: Substrate sequences by cleavage site.

As noted when comparing MPro structures, the N142C change in MPro2 and MPro3 might be expected to reduce stringency for Q at P1, and indeed Cluster 2 has the lowest frequency (though still 97%) of Q, and Cluster 3 is not uniformly Q either, though the N142A change in MPro5 did not produce a corresponding reduction. The P168N exchange in MPro4 may form hydrogen bonds with P3 and/or P5 amino acids, possibly explaining the fact that Cluster 4 is the only example wherein lysine is the most prevalent at P5 and somewhat common at P3. (N.B.: Lysine is negligible in other clusters at P5 and far less common in P3 except for Cluster 2.)

Similar to active site residues, we separated substrate sites by cluster for distance comparison (Figure 6B). Comparing Figure 6B to Figure 5C, we see that the rank-order of clusters from the cluster center is not the same, meaning that conservativity between active site and substrate is not a perfect correlation. However, some patterns are retained; for example, Cluster 1’s closest neighbor, as expected from the cladogram, is Cluster 2 for both active site and substrate sequences.

Within a given polyprotein sequence, the 11 cut sites are not identical sequences, thus we also performed a distance comparison of sequences divided by cut site (Figure 6C). Despite larger sample sizes, the distances between frequency matrices distinguished by cluster are smaller than those distinguished by cleavage site, suggesting that diversity within the polyprotein sequences of similar species exceeds cross-species diversity, further supporting the possibility of cross-compatibility between different coronaviral MPro enzymes.

### Experimental protease substrate validation

Predicted substrates for the representative MPro enzymes were screened in a yeast-based protease activity reporter called YESS 2.0^15^ and analyzed by flow cytometry to validate predictive model accuracy. In YESS 2.0, the protease and substrate cassettes are encoded on a single plasmid under the control of β-estradiol and galactose-inducible promoters, respectively. The substrate polypeptide contains an AGA2, followed by a FLAG tag, a substrate sequence, an HA tag, and a WEHDEL ER retention signal. The protease is preceded by an ER targeting signal and succeeded by a strong WEHDEL ER retention signal^16^ (Figure 7A). Upon induction with β-estradiol and galactose, the protease and substrate polypeptides are trafficked to the ER. As the polypeptides travel through the ER, the encoded substrate, flanked by FLAG and HA tags, can be cleaved by the protease. On the cell surface, a fully intact substrate cassette, resulting from an uninduced or inactive protease, will retain both the FLAG and HA tags. Upon cleavage, the substrate will lose the HA tag but retain its FLAG tag. The epitope tags can be stained with fluorescently labeled antibodies, resulting in a high fluorescence signal of anti-FLAG and anti-HA fluorochromes for an intact substrate and a low HA/high FLAG signal for cleaved substrates (Figure 7B). Screening of predicted substrate sequences for MPro1 (alphacoronavirus), MPro2 (murine coronavirus), MPro3 (duck coronavirus), MPro4 (quail coronavirus), and MPro5 (avian coronavirus) were performed in this fashion (see Methods), to determine active protease-substrate pairings for each enzyme target. Protease activity was quantified by calculating the fold change in normalized fluorescence signal (anti-Flag-PE/anti-HA-Alexa 647) between protease-induced and uninduced samples (Figure 7C).

**Figure 7.**
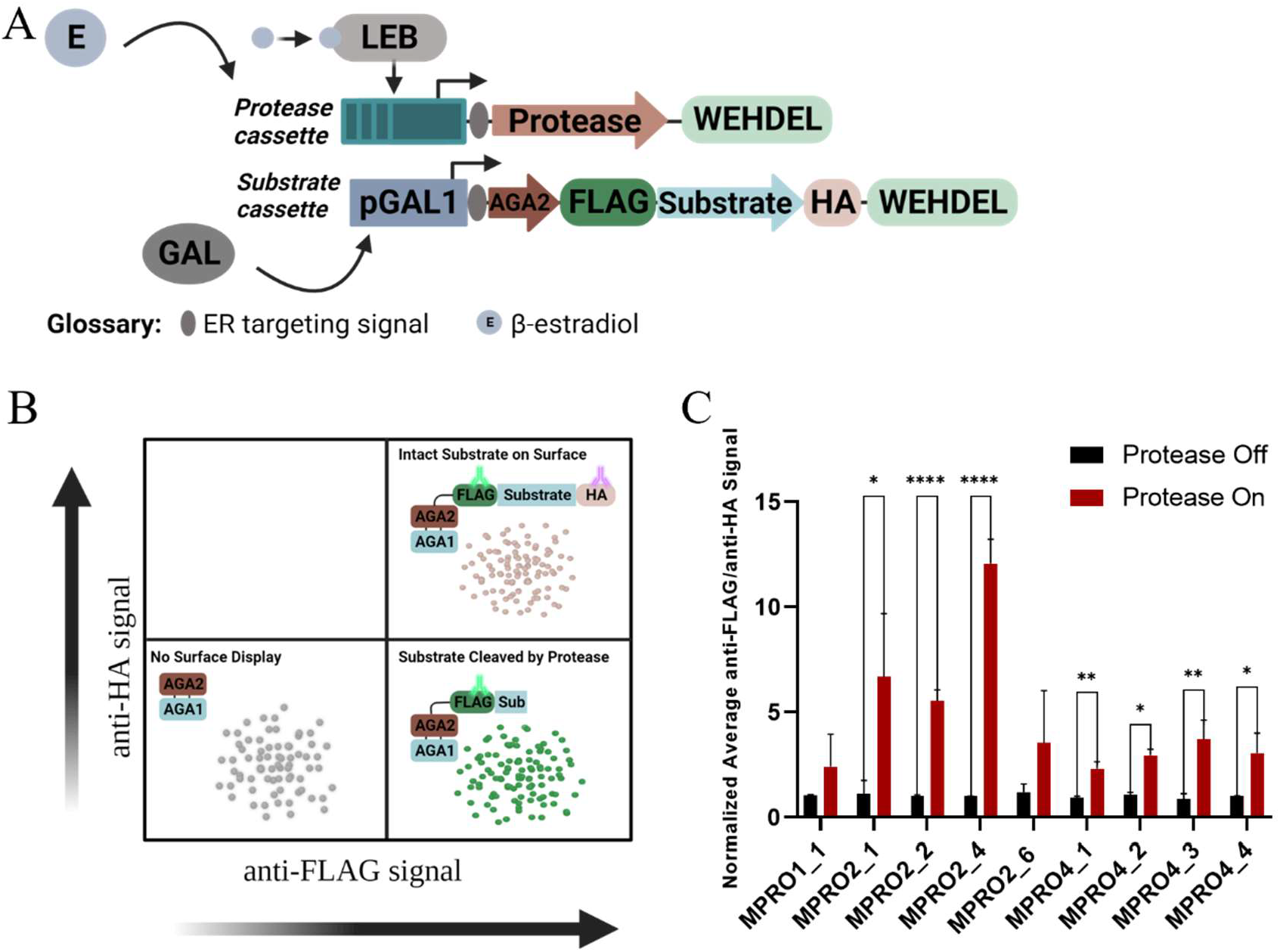
A: The architecture of the protease and substrate cassettes. B: The expected phenotypes from protease activity on the yeast cell surface. C: Signal ratios of cleaving proteases.

Three of the five representative MPro enzymes were active on at least one of their predicted substrates (Figure 7C). A single substrate for MPro1 (IKVSTIQSKLT) was tested in the system and was observed to be an active protease-substrate configuration, with a 2.5-fold increase in activity when compared to control. MPro2 was active on four of six predicted substrates (LAGVKLQSKRT, IEVSQIQSRLT, VSTVVLQNNEL, RDNTVLQALQS), with activity fold changes between 3-fold to 12-fold when compared to the uninduced protease (see Methods). MPro4 was also active on four of the six predicted substrates (EHKTVVQAVAD, IAVSTVQNKIL, LTFTNLQNLEN, SGTTILQAGTH), with activity fold changes between 2-fold to 4-fold when compared to the inactivated protease cassette of the uninduced samples (see Methods). For unclear reasons, all six substrates tested against MPro5 resulted in no significant change in mean fluorescence signal ratios. It is possible that MPro5 is inactive in our assay. Similarly, MPro3 showed no activity on the single tested substrate, with a sequence inferred from the protease termini since the NCBI sequence included only the protease. Nonetheless, this yeast platform remains a rapid method to confirm and measure protease activity and shows that identified substrates for 3 out of 5 proteases are indeed cleaved as anticipated from computational analyses.

### MPro inhibition by nirmatrelvir

Pfizer recently discovered and developed an oral MPro inhibitor, PF-07321332^8,11^ or nirmatrelvir. In *in vitro* energetic assays, the inhibitor was effective across multiple coronavirus species, including both alphacoronaviruses and betacoronaviruses (including SARS, MERS, and SARS-CoV-2). Given that the inhibitor was effective against multiple branches of the phylogenetic tree, we sought to evaluate its efficacy more broadly against our representative MPros. We developed an inhibition assay, based on activity change on successfully cleaved substrates in YESS 2.0 described above.

First, we sought to replicate SARS-Cov2 MPro inhibition by nirmatrelvir in our system. In this case, the inhibition assay was conducted similarly to the activity assay, except that nirmatrelvir was added to the culture. Just as the activity of a protease on its target substrate in this system is observed with a loss of the HA epitope tag, indicated by an increased ratio of the normalized fluorescence signal (anti-Flag-PE/anti-HA-Alexa 647), the inverse can be quantified for an inhibited protease. For consistency, the Protease On and the Protease On + Drug sample sets were normalized to the Protease Off sample set (Figure 8A). This normalization allows for a direct comparison of activity and inhibition, with a ratio of >1 signifying activity and a decrease in that signal representing inhibition.

**Figure 8.**
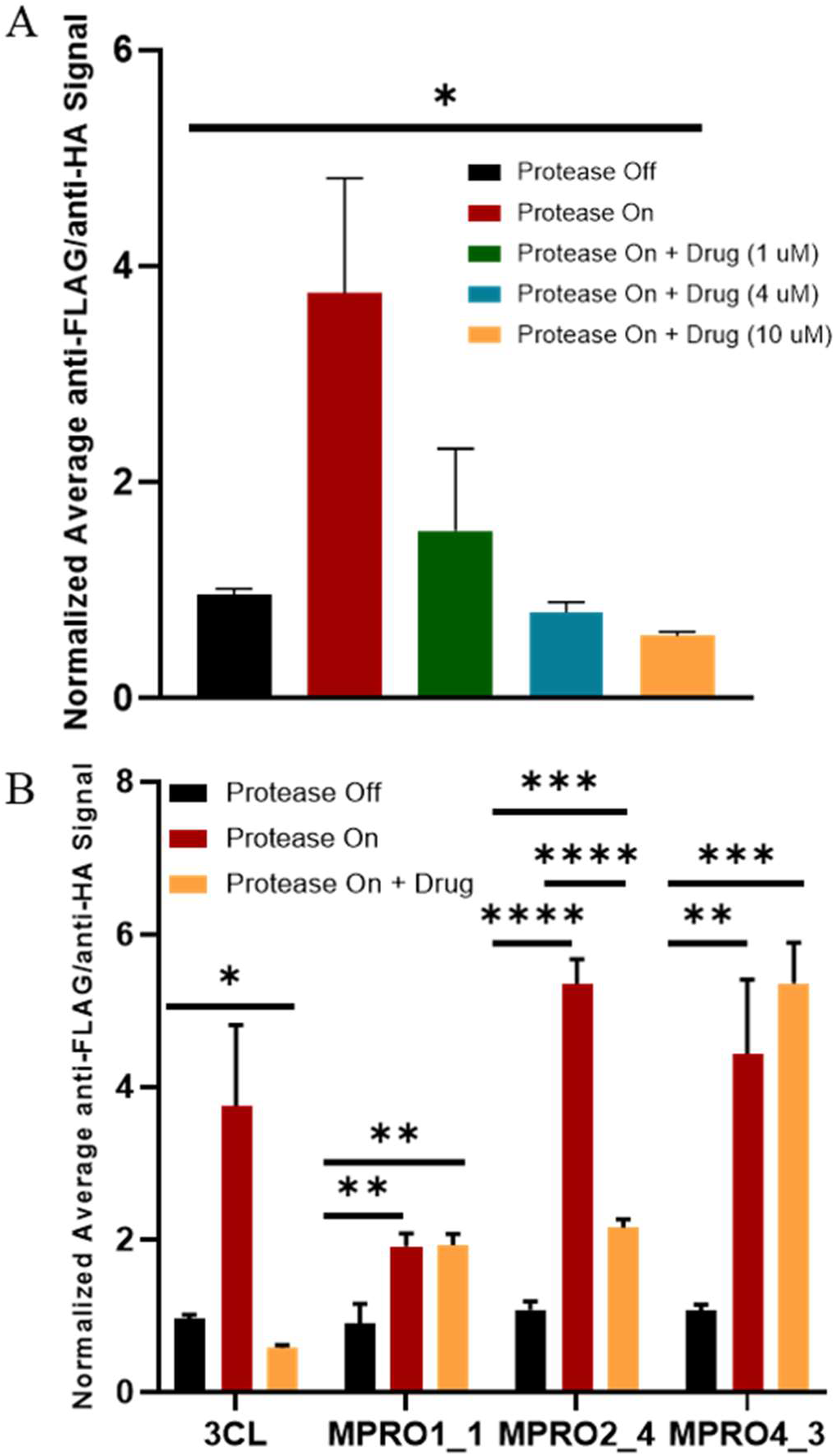
Yeast surface display-based cleavage assays. A: Titrating nirmatrelvir inhibition of SARS-CoV2 MPro. B: Nirmatrelvir inhibition of coronavirus main proteases (MPRO) of different species origins. Substrate identities are as follows: MPro1 (1 – IKVSTIQSKLT), MPro2 (4 – VSTVVLQNNEL), MPro4 (3 – LTFTNLQNLEN). The anti-FLAG-PE/anti-HA-Alexa 647 fluorescence ratio observed when protease expression is induced is divided by the normalized anti-FLAG-PE/anti-HA-Alexa 647 fluorescence ratio in the absence of protease. The baseline fluorescence in the absence of protease activity is defined as approximately 1, and values ≤1 are considered no activity. A normalized ratio >1 corresponds with protease activity. *p ≤0.05, **p ≤0.01, ***p ≤0.001, ****p ≤0.0001. n=2 3CL, n=3 MPro.

We find that SARS-Cov2 MPro inhibition was concentration dependent (Figure 8A). While 50% inhibition is observed at 1 μM nirmatrelvir, SARS-Cov2 MPro is completely inhibited in the presence of 4 and 10 μM. A 10 μM nirmatrelvir concentration was chosen to screen inhibition of MPro from the different species. From the experimental validation of predicted substrates, the most active protease-substrate pairs were selected for this inhibition assay; MPro1-1 (IKVSTIQSKLT), MPro2-4 (VSTVVLQNNEL), MPro4-3 (LTFTNLQNLEN) (Figure 8B). From the three protease-substrate pairs tested for nirmatrelvir-induced inhibition, MPro2 activity on its target substrate VSTVVLQNNEL was the only pair to show significant inhibition, a 4-fold decrease in activity, in the presence of 10 μM nirmatrelvir (Figure 8B). Nirmatrelvir did not significantly inhibit MPro1 or MPro4, which retained full activity in the presence of 10 μM nirmatrelvir (Figure 8B).

## Discussion

Preparedness against future pandemics caused by coronaviruses requires the development of an arsenal of inhibitors against the broad diversity of coronaviral targets such as MPro enzymes. In this study, we identified and analyzed the structures and sequences of several representative MPro sequences for all sequenced coronaviral species. We experimentally validated cleavage of substrates identified from polyprotein sequences and evaluated inhibition by nirmatrelvir. We demonstrated that nirmatrelvir was effective against Mpro2 (murine coronavirus) activity on its highly active substrate (VSTVVLQNNEL) with a similar degree of inhibition as SARs-CoV-2 MPro but was ineffective against two other identified homologous MPros (MPro1 and MPro4). Further sampling from the cluster centers in the active site cladogram can be used to identify the breadth of nirmatrelvir’s effect, and by exclusion, identify which coronaviral species are not effectively inhibited by this inhibitor. In this manner, we provide a method for evaluating all future MPro inhibitors for the breadth of their inhibition.

The inactivity of nirmatrelvir against MPro1 and MPro4 underscores the importance of this study–while nirmatrelvir was demonstrated to be an effective *in vitro* inhibitor of several alpha- and betacoronaviruses (Figure S1), our study suggests that additional inhibitors will need to be evaluated to achieve pan-coronaviral preparedness. The active site similarity matrix (Figure 5C) suggests that the best inter-cluster cross-inhibition between Clusters 3 and 5, and the worst between 1 and 5, and 2 and 4. This reasoning would reduce our expectation that a member of Cluster 1 would be inhibited by an inhibitor developed for Cluster 2 (such as nirmatrelvir), though it would be expected to have higher likelihood of inhibition than a member of Cluster 4. Cluster 1 notably exhibited high active site sequence variability in comparison to other clusters (Figure 5B), which may help to explain why MPro1 is not inhibited in our study, while some other members of Cluster 1 were found to be potently inhibited by nirmatrelavir^8^. Structural modeling of nirmatrelvir binding to AF2 models of MPros (structures provided in SI) does not offer any obvious explanation for the observed differences in inhibition (Table S11), suggesting that further experimental characterization, including using MX, may be necessary to shed light on the underlying molecular mechanisms. While promising, AF2 models clearly remain an insufficient sole basis for inhibitor development.

A notable limitation in our study is that we are examining diversity across only MPro representatives with known polyprotein sequences. It is likely that many more coronaviral species exist for which genome sequences are not available. Moreover, these MPros have evolved in the absence of selection pressure from protease inhibitors. As we use nirmatrelvir and other inhibitors to respond to the current pandemic, it is likely that escape mutants will develop^11,17–19^. Escape mutations are typically deleterious and therefore uncommon in the absence of inhibitors, and therefore are likely absent from our dataset. Further studies will need to screen for potential escape mutants in the identified MPros, and similarly proactively develop inhibitors that will be effective against them.

The failure to use our knowledge of SARS, MERS, and coronaviruses broadly is one of many reasons that national and regional public health systems were unprepared to mitigate the spread of SARS-CoV-2, resulting in 15 million excess deaths^20^ and $16 trillion in economic damage^21^. Had we, as a scientific community, prepared to respond to future coronaviral threats when beset by SARS and MERS, it is likely that the costs of the SARS-CoV-2 pandemic in lives and economic damages could have been substantially mitigated. Our intention in this work is to help lay the groundwork for avoiding repetition of that costly failure to prepare by looking ahead to future potential pathogens. It is almost certainly only a matter of time before another species of coronavirus is able to infect and spread within human populations and, given the incidence of co-infection and high rates of recombination between coronaviruses, it may not be another betacoronavirus. Our work will aid in determining whether a safe and effective inhibitor of the SARS-CoV-2 main protease may be effective against other coronaviruses, or if additional drugs might need to be discovered and developed. For future anti-coronaviral agents, it would be ideal to not only screen against the currently prevalent viral species drug targets, but also against representative homologs across all coronaviral genera. We further suggest that if some developed inhibitors fail to be effective against other MPros, that other inhibitors be proactively developed and tested to ensure that we have defenses ready against the next coronaviral agent that emerges, whatever it may be.

## Methods

### Reference proteases

The SARS-CoV-2 MPro reference 3D structure used in this study was PDB ID 6yb7^22^. We also compare with other MPro CSMs with other publicly available experimentally-determined structures in the Protein Data Bank^23–25^: SARS (1q2w^26^ and 1wof^27^), MERS (4ylu^28^), PEDV (6l70^29^), TGEV (2amp^27^), HKU4 (2yna^30^), A59 (6jij^31^), HKU1 (3d23^32^), FIPV (5eu8^33^), NL63 (3tlo^34^).

### Sequence alignment for database assembly and filtering

Sequence alignments for database assembly and filtering were performed with the BioPython^35^ globalms function, using an identity reward of 1, a mismatch penalty of −0.2, an open gap penalty of −2, and an extend gap penalty of −0.1, with no penalty on end gaps. Single best alignments were used.

### Database assembly and filtering

We assembled a list of 346 MPro sequences with <99% sequence identity as follows. We collected a list of protein and polyprotein sequences from the NCBI Virus database^10^ by filtering the sequences to coronaviridae (taxid 11118). Downloaded 6/11/2021, this list included 4,266,081 members. We excluded all sequences with species listed as either ‘Severe acute respiratory syndrome-related coronavirus’ or ‘Middle East respiratory syndrome-related coronavirus’ as these proteases were highly similar and reduced the list to 53,623. Database sequences were aligned with the SARS-CoV-2 MPro reference sequence, and where the alignment score exceeded 10, the region of sequence that matched the reference was collected, yielding a set of 3,278 MPro sequences. An identity matrix for these sequences was generated using Clustal Omega^36^ and all members with identities ≥ 0.99 below the diagonal were excluded, yielding 346 MPro sequences.

### Computed structure model generation

All 346 MPro sequences were modeled via Robetta (https://robetta.bakerlab.org/), using the trRosetta algorithmm^37^. Robetta produced five candidate CSMs, and the CSM with the lowest RMSD to any comparison PDB experimental structure was used for selection of representative structures. Upon public availability, we used AlphaFold2^38,39^ to henerate higher-quality CSMs. For each sequence, we generated five monomeric structure models and used the one with the highest pLDDT score. We repeated this protocol modeling each MPro as a dimer.

A comparison of CSMs showing alignment lengths and RMSDs between SARS-CoV-2 MPro, the CSMs used for representative selection, the AF2 monomer, and both chains of the AF2 dimer is in Table S12. AF2 CSMs were highly similar to those generated using trRosetta, with only four with RMSD > 1.5 Å. (N.B.: In two of those cases, there was also disagreement between AF2 monomer and dimer structures > 1.5 Å, suggesting uncertainty in modeling those sequences with either method.) Because of the similarity, we considered the selections we made based on trRosetta models to be sufficiently representative without repeating the experiments.

To generate substrate-bound models, an initial peptide structure was generated for the native MPro nsp4-nsp5 cleavage site (ITSAVLQ/SGFR) and superimposed over the inhibitor of 6lu7^40^. The peptide was then positioned using the Rosetta FlexPepDock protocol^41^. Substitutions of the substrate sequence to convert into each of the native cleavage substrates was done using the Rosetta FastRelax protocol^42^. For both Rosetta protocols, geometric constraints were used to preserve the catalytic geometry of H41 and C145, as well as the P1 residue relative to C145 and the oxyanion hole. Similar procedures were used to generate bound structures of the representative MPro complexes, using the AlphaFold structure instead of 6lu7. For nirmatrelvir-bound models, only the FastRelax protocol was used, adding distance constraints to enforce the crystal structure’s H-bonds.

### Active site identification and alignment

The CSMs were aligned for active site identification using Dali^12^. Active site residues of the reference proteases were identified as those with any atom within 5.5 Å, or with C_α_ within 9 Å of the substrate and C_α_-C_β_ vector within 75° of a substrate atom, yielding the following residues: 24, 25, 26, 27, 39, 41, 49, 54, 140, 141, 142, 143, 144, 163, 164, 165, 166, 167, 168, 172, 187, 188, 189, 190, 191, and 192. With structurebased sequence alignment, active site residues were identified in all other models as those which aligned with a reference MPro active site residue.

### Cladogram generation and selection of representative models

The full protease and active site cladograms were generated using MUSCLE^43^. The active site cladograms were used to select representatives. The sequences tended to group by genera (alpha, beta, gamma, and delta), and so genera-based cluster representatives were selected. We generated an all-against-all structural Z-score matrix for all models in the cluster using Dali^12^ and selected the model with the highest sum of all non-self Z-scores as the cluster representative.

### Protease specificity identification

To identify the canonical cleavage sites, we first performed sequence alignment of the NCBI entry sequences of all accession numbers in MPro lists using MAFFT^44^. Cleavage sites were identified for SARS-CoV-2 and aligned sites in other sequences were extracted as the homologous cleavage site. Some sequences were single proteins, so no cleavage sites could be identified in this manner. Aligned sites with deletions/exclusions compared to the reference were also not considered valid. Consequently, our analysis was not able to include 11 canonical cleavage sites for all enzymes. This was the case for the Cluster 3 representative, so its modeled substrate was an inference based on the protease’s own termini.

### Substrate and protease cloning in pY2 plasmid

To determine active substrates for MPro targets, a high-throughput analysis of protease activity was conducted. From the list of predicted substrates for each of the protease targets (Table S3), oligo pairs for each substrate were designed as Part 2 for the substrate cassette of the YESS protease-substrate expression system^15^ (Figure 7A). Oligonucleotides were annealed and phosphorylated with T4 PNK and diluted to a final concentration of 3 μM in water. Additional oligonucleotides were also prepared to assemble the substrate cassette – FLAG epitope tag (Part 1), HA epitope tag (Part 3), WEHDEL ERS (Part 4). The substrate cassette was constructed into the YESS plasmid for each substrate target using a *BsmbI* Golden Gate reaction^45^. The assembled plasmid was transformed into competent *E. coli* (ZymoResearch T3001) cells and plated on selection media (Ampicillin, Goldbio 69-52-3). A green-white screen^46^ was used to inoculate assembled colonies in selection liquid media. Correctly assembled plasmids were confirmed by Sanger sequencing^47^ (Genewiz, Azenta Life Sciences). Assembled YESS plasmids with correct substrates were used as the backbone for a second Golden Gate reaction, using *BsaI*, to assemble the protease cassette. Proteases of interest were purchased as gene fragments from Genewiz and contained the WEHDEL ERS and correct overhangs. Once completed, the reaction mixture was transformed into competent *E. coli* (ZymoResearch T3001) cells and plated on selection media (Ampicillin, Goldbio 69-52-3). Sanger sequencing confirmed plasmids that contained the correct protease insert. MPro5 substrates SNVVVLQSGHE and KSFSALQSIDN did not anneal during design and were omitted from the assay, since six other substrates were considered sufficient.

### Flow cytometry analysis of protease activity

YESS plasmids harboring substrate and protease cassettes were transformed into an engineered EBY100 yeast strain expressing a LexA-hER-haB112 transcription factor necessary for β-estradiol-induced expression. Transformed yeast cells were grown in a YNB-CAA-glucose medium for 24 hours at 30°C in YNB-CAA, 2% Glucose, 2% Raffinose. Saturated cultures were inoculated to an OD600 of 1 in 0.25 mL of YNB CAA Glucose Raffinose, 2% Raffinose, and grown to an OD_600_ range of 2-4 at 30°C in a deep-well 96-well plate (ThermoFisher Scientific, 260252 https://www.thermofisher.com/order/catalog/product/260252). Once reached, the outgrow OD_600_ was used to calculate the amount of culture required to induce the samples at a starting OD_600_ of 0.5 in 0.25 mL media. Required culture volumes, two per original sample, were washed in minimal media (YNB CAA) with galactose as the carbon source to remove residual glucose from the cells. Supernatant from the wash was removed and the cells were then induced for protein expression. One sample set (Protease Off) was induced in YNB CAA Galactose to induce substrate expression and the other (Protease On) was induced in YNB CAA Galactose and β-estradiol (2 μM) to induce substrate and protease expression. The dual-induction allows for an observable fold-change difference because of protease activity when the enzyme’s expression is induced. Once all cultures were induced to appropriate starting OD_600_ for growth, the plate (deep-well 96-well plate) was set to shake at 30°C for 12-16 hours. Grown induction culture was analyzed the next day to calculate the OD_600_ for each sample, with resulting values used to determine the culture amount equivalent to approximately two million cells. Each sample of two million cells were washed in a round-bottom 96-well plate (ThermoFisher Scientific, 174929) in phosphate-buffered saline (PBS) with bovine serum albumin (BSA) (0.5% BSA, Goldbio 9048-46-8). Washed cells were stained with two fluorescently labeled epitope tag-specific antibodies (anti-FLAG PE, Biolegend, cat# 637309), anti-HA Alexa 647, cat #682404) and incubated at room temperature in the dark for 90 minutes. Stained cells were washed with PBS BSA (0.5% BSA) and resuspended in 0.2 mL PBS BSA (0.5% BSA). Stained cells were assayed by flow cytometry (NL Cytek 3000, Cytek Biosciences). Protease Off samples were used as controls to determine the display of the intact (non-cleaved) substrate cassette on the surface of the cells, as these samples have both epitope tags displaying. Analysis of protease activity is based on the removal of the C-terminal HA epitope tag because of the substrate sequence being cleaved by the active protease. Comparative analysis was conducted to assay fold-change differences in anti-HA display signal between the uninduced and induced proteases to determine valid substrate targets within yeast for the corresponding protease.

### Testing Inhibition

YESS plasmids harboring the selected MPro protease-substrate pairs producing the clearest activity signals were transformed into an EBY100 yeast strain and the outgrow protocol followed as described above. Cells were washed in minimal media (YNB CAA) with galactose to remove residual glucose media before three sample groups were induced. The first sample set (Protease Off) was induced with YNB CAA Galactose to induce expression of the substrate cassette only. The second sample set (Protease On) was induced with YNB CAA Galactose and β-estradiol (2 μM) for expression of both the protease and substrate cassettes. The third sample set (Protease On + Drug) was induced with YNB CAA Galactose, β-estradiol (2 μM), and nirmatrelvir (10 μM). Induction time and temperature, as well as staining protocols, were followed as described above. All three sample sets were assayed by flow cytometry (NL Cytek 3000, Cytek Biosciences).

## Supporting information

Supplementary tables, figure, and molecular structures

## Acknowledgements

RCSB PDB members are supported in these activities by the National Science Foundation (DBI-1832184), the US Department of Energy (DE-SC0019749), and the National Cancer Institute, National Institute of Allergy and Infectious Diseases, and National Institute of General Medical Sciences (NIGMS) of the National Institutes of Health (NIH) under grant R01 GM133198. We gratefully acknowledge support the Rutgers University RISE (Research Intensive Summer Experience) Program for Cassandra Olivas (and NSF-REU), Mickayla Bacorn (and UMBC U-RISE NIGMS/NIH T34 GM 136497), MaryAgnes Balogun (and Aman Armaan Ahmed Family), Amy Wu-Wu (NSF-REU). Lastly, we acknowledge start-up funding from the University of Florida Chemical Engineering Department, and funding from the National Institutes of Health (NIH) under grants R21GM144812 and R35GM146821 (Carl Denard, Sam Martinusen).

## Notes

### Competing Interest Statement

The authors have declared no competing interest.

